# Single-cell colocalization analysis using a deep generative model

**DOI:** 10.1101/2022.04.10.487815

**Authors:** Yasuhiro Kojima, Shinji Mii, Shuto Hayashi, Haruka Hirose, Masato Ishikawa, Masashi Akiyama, Atsushi Enomoto, Teppei Shimamura

## Abstract

Analyzing colocalization of single cells with heterogeneous molecular phenotypes is essential for understanding cell-cell interactions, cellular responses to external stimuli, and their biological functions in diseases and tissues. However, high-throughput methods for identifying spatial proximity at single-cell resolution are practically unavailable. Here, we introduce DeepCOLOR, a computational framework based on a deep generative model that recovers inter-cellular colocalization networks with single cell resolution by the integration of single cell and spatial transcriptomes. It segregates cell populations defined by the colocalization relationships and predicts cell-cell interactions between colocalized single cells. DeepCOLOR could identify plausible cell-cell interaction candidates in mouse brain tissues, human squamous cell carcinoma samples, and human lung tissues infected with SARS-CoV-2 by reconstructing spatial colocalization maps at single-cell resolution. DeepCOLOR is typically applicable to studying cell-cell interactions in any spatial niche. Our newly developed computational framework could help uncover molecular pathways across single cells connected with colocalization networks.

## 2 Introduction

Single-cell analysis of transcriptome profiles has given rise to novel avenues for examining heterogeneous cell populations. Such techniques demonstrate continuous cell states rather than classical discrete cell types [22, 43]. The heterogeneous cell states observed via these techniques were shown to be crucial for elucidating mechanisms underlying disease development and prognosis [14, 21]. Hence, it is important to understand the molecular mechanism involved in the transition between the cell states and their regulation. This heterogeneity can be attributed to the environmental cues obtained from surrounding cells [30], which is termed as cell-cell interaction (CCI). Environmental cues facilitating CCI rely on heterogeneous molecular states of the surrounding cells (i.e. the expression of ligands and the stiffness of cells). Hence, to dissect the molecular basis of CCI, it is crucial to analyze colocalization patterns among single cells with heterogeneous molecular phenotypes. However, the observed single-cell transcriptome loses the spatial context when analyzed using the available high-throughput methodology.

Recently, spatial transcriptome observation technologies, such as spatial transcriptomics (ST) [44], Visium (10X Genomics), Slide-seq [41], and high-definition spatial transcriptomics [46], have enabled analysis of the entire transcriptome spatially. These technologies provide a unique opportunity to characterize local niches with comprehensive molecular profiles. For example, Ji et al. utilized this technology to show that the unique population in squamous cell carcinoma is resident in the leading edges of the tumor [25]. In contrast, widely used high-throughput methods of spatial transcriptomics such as ST and Visium have limited capture rates (resulting in substantial spatial dropout that increases with higher resolution) and do not achieve single-cell resolution. In situ sequencing, MERFISH [49], seqFISH [17], and other in situ techniques allow spatial analysis at single-cell resolution. However, these techniques require high expertise in experimental technologies and capture relatively fewer pre-specified genes than the sequence-based methodologies. These inherent limitations of current spatial transcriptome analysis techniques prevent the identification of interaction networks between heterogeneous cell populations involved in disease progression.

Several computational methodologies have been developed to address this limitation by the integration with scRNA-seq observation. One major approach involves deconvolution of observations at each spot of the spatial transcriptome with a cell-type expression profile [1, 6, 16, 29]. This approach reveals the spatial distribution of heterogeneous cell types, which have been identified in the scRNA-seq analysis. In contrast, cell type-based analyses have a limitation in that they can analyze colocalization only between predefined populations, which is highly dependent on the manually determined clustering resolution and does not necessarily correspond to the differences in the spatial distribution of the tissues. While a recent single-cell approach successfully revealed cell distribution across various spatial contexts [4], the methodology was not suitable for analyzing the colocalization network as it could estimate distinct spatial assignments among cells with almost the same profiles. Hence, a new computational approach is required for identifying the colocalization between heterogeneous cell states captured by scRNA-seq.

DeepCOLOR is a deep learning framework for addressing the dual challenges of (1) recovering spatial contexts of single cells observed in scRNA-seq data and (2) revealing cell-cell interactions in spatial niches. DeepCOLOR was used to build a continuous neural network map from latent cell state space to each spot in the spatial transcriptome in order to enhance consistent mapping profiles between single cells with similar molecular profiles. This mapping technique did not rely on the fine-grained cell type preparation, which is the upper limit of the resolution for many of the existing deconvolution methods [1, 6, 16, 29], and revealed the spatial distribution of all single cells observed by scRNA-seq with high accuracy. A colocalization profile between two cells could be derived from the overlaps between the spatial distributions of individual cells, and clustering analysis of neighboring single-cell pairs could classify colocalized cell populations beyond the resolution at the cell type level. In addition, DeepCOLOR demonstrated the ability to predict ligand-mediated cell-to-cell communication by combining colocalization scores between cells, gene expression within cells, and prior knowledge of signal transduction and gene regulatory networks. These single cell colocalization-based analysis enabled us to extract and characterize the population which affected by environmental ques from surrounding cells with unprecedented specifity, which was potentially overlooked by predefined population-based colocalization analysis [2, 36, 37]. We used DeepCOLOR to reveal intercellular communication mechanisms in two complex tissues. This study could provide valuable insights regarding the application of DeepCOLOR as an analytical tool for studying cell-cell interactions in any spatial niche.

## 3 Results

### 3.1 Methodology overview

We developed DeepCOLOR, a deep generative model for colocalization representation, which uses scRNA-seq data as a reference cell population to deconvolve each spot of the spatial data and decipher the intercellular colocalization network across single cells in a spatial niche. The input to DeepCOLOR was scRNA-seq data along with spatial transcriptome data derived from the same tissue or region discerned based on currently available spot-wise spatial methods (Visium, ST, GeoMx, etc.). The following assumptions were made for the input: the two modalities share some subset of common genes, and the variations in the single-cell transcriptome cover the cell populations within the spatial niche. DeepCOLOR first transformed the transcriptome observation of each cell from scRNA-seq data into a latent cell state via encoding by a pre-trained neural network. It then optimized an objective function to maximize the likelihood for the probability model of the spatially distributed transcriptome observed in the spatial transcriptome data. DeepCOLOR then provided spatial distributions of each cell in the scRNA-seq data, which comprised information about singlecell localization, indicating the likelihood of the presence of all cells in the scRNA-seq data at each spot in the spatial transcriptome. This information enabled us to search for colocalized pairs of cells defined by the overlaps of spatial distributions. With its ability to smoothly map the learned latent representation of a single cell to a spatial spot, DeepCOLOR could (1) correct poor-quality spatial measurements attributed to technical variability such as spatial dropout; (2) estimate spatial distributions at the single-cell level independent of predefined labels such as cell types; (3) identify and classify cell colocalization pairs that exist in spatial niches; (4) identify differentially expressed genes in colocalized cell clusters; and (5) identify ligands involved in cellular communication between colocalized cell population.

Technically, DeepCOLOR was based on latent representations of single cells derived from deep generative models and optimization via a stochastic gradient descent method. Unlike the conventional application of deep generative models to scRNA-seq data [18, 33], the optimization function involved the likelihood of the probabilistic model of gene expression in each spatial spot, which is assumed to be the weighted average of the transcriptome associated with latent cell states in the scRNA-seq dataset. DeepCOLOR modeled the difference in the technical capture rate for each gene when comparing scRNA-seq data and spatial transcriptome data and the abundance of each gene in the spatial transcriptome assuming correction terms for the sensitivity and contamination of technical measurements. DeepCOLOR could then estimate a continuous mapping function that inferred the spatial distribution of single cells based on their latent cell states, thus obtaining a consistent spatial distribution among single cells with similar molecular profiles. To further strengthen the consistency of the mapping function, we employed a stochastic gradient descent method using mini-batches of single cells. This formulation imposed a constraint: a subset of single cells obtained by down-sampling forms consistent spatial expression patterns, and hence cells with approximately the same latent representation would map to the same spatial location. Downstream analysis of DeepCOLOR implemented single-cell colocalization clustering based on latent representations of cell pairs that were likely to colocalize, differential expression analysis in specific colocalized clusters based on the Willcox rank-sum test, and ligand activity analysis between colocalized cell populations. DeepCOLOR has been implemented as a PyTorch module and is available at https://www.github.com/kojikoji/deepcolor.

### 3.2 Accurate mapping with well-known spatial domains

To validate our newly developed assignment methodology, we applied our mapping algorithm to a mouse visual cortex dataset [29]. First, we evaluated the similarity of the aggregated gene expression of single cells assigned to each spot with the original gene expression of the spot. We found that the expression was well correlated for genes with high expression even if the genes were not used for training of the generative model (**Fig. 1-a in Supplementary**). Next, we validated whether the mapping patterns of single cells were consistent with the well-known anatomical structures. We found that the aggregated assignment of the excitatory neuronal subtypes associated with L2/L3 layers (Ext L23) was enriched in the outer region compared to that of subtypes associated with L5/L6 layers (Ext L5 1 and Ext L56) (**Fig. 2-a, b**). In contrast, single cells within each layer were assigned to more specific regions compared with the aggregated spatial distribution of the predefined subtypes described above (**Fig. 2-a, c**). These results suggested that DeepCOLOR captured the molecular signatures of single cells obscured by clustering analysis and utilized them for reconstructing the spatial distribution of single cells. We also found that the assignments of single cells to a specific spot performed by DeepCOLOR were smooth on the latent cell state space of single cells (**Fig. 2 in Supplementary**). This smooth assignment enabled us to capture the colocalization between single cells as the overlap of the estimated spatial distribution. We examined the colocalization scores for single-cell pairs across the excitatory neuronal subtypes associated with cortical layers to validate our approach to extract colocalized single-cell pairs. We found that a larger proportion of single-cell pairs across expected adjacent layers (among L2 and L3 or among L5 and L6) demonstrated high colocalization scores (*c*_*ij*_ *>* 1)than those across more distant layers (**Fig. 2-e**). These results showed that our colocalization scoring methodologies were useful for identifying the adjacent cell population for every single cell.

**Figure 1:**
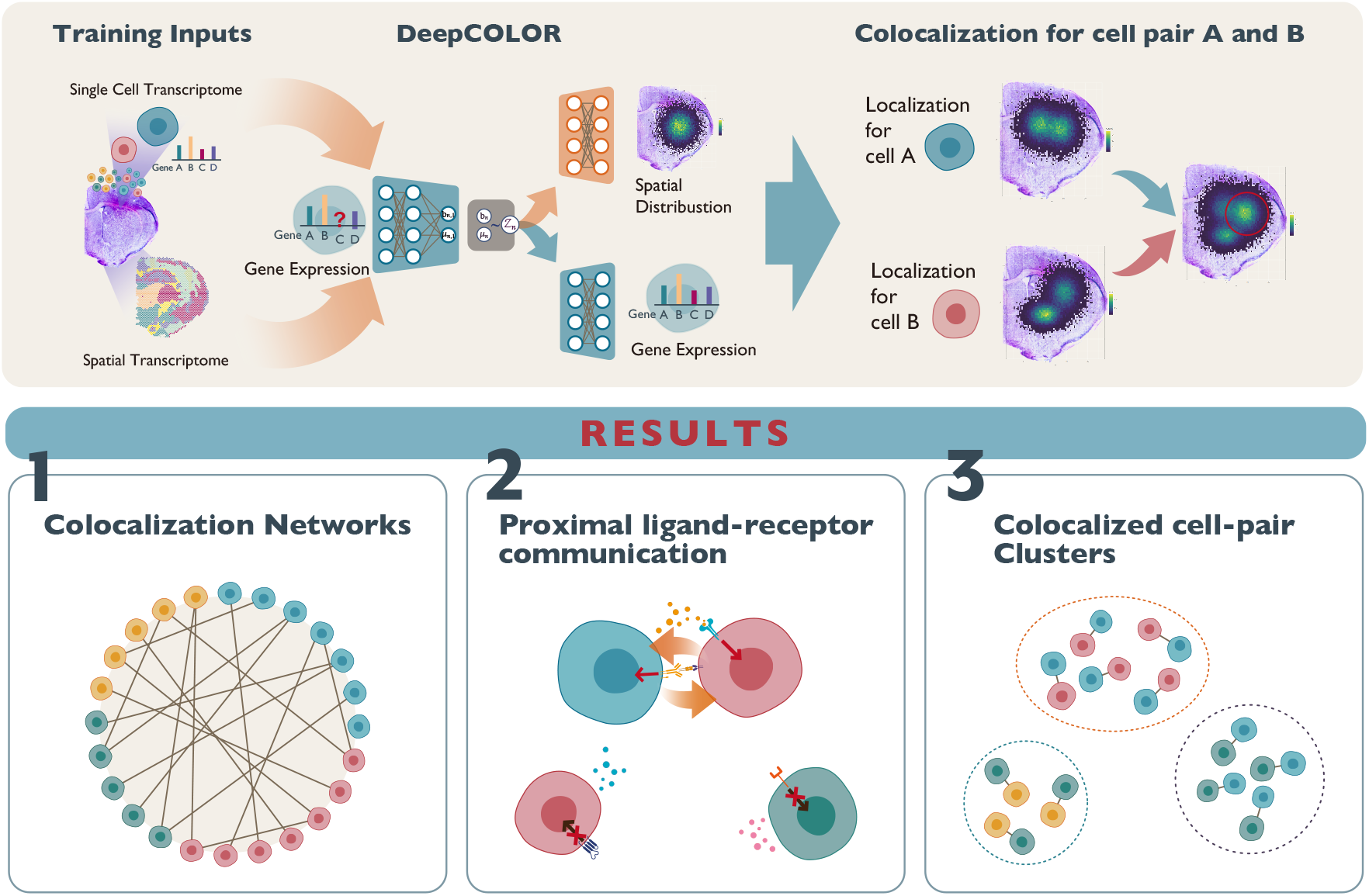
Schematic representation of the workflow of DeepCOLOR. DeepCOLOR takes single cell and spatial transcriptome as traning inputs and reconstruct spatial distribution and denoised expression profile from noisy single cell observation. Using spatial distribution, we can evaluate colocalization relationships between single cells and identify colocalization network, proximal ligand-receptor communication and colocalized cell-pair clusters.

**Figure 2:**
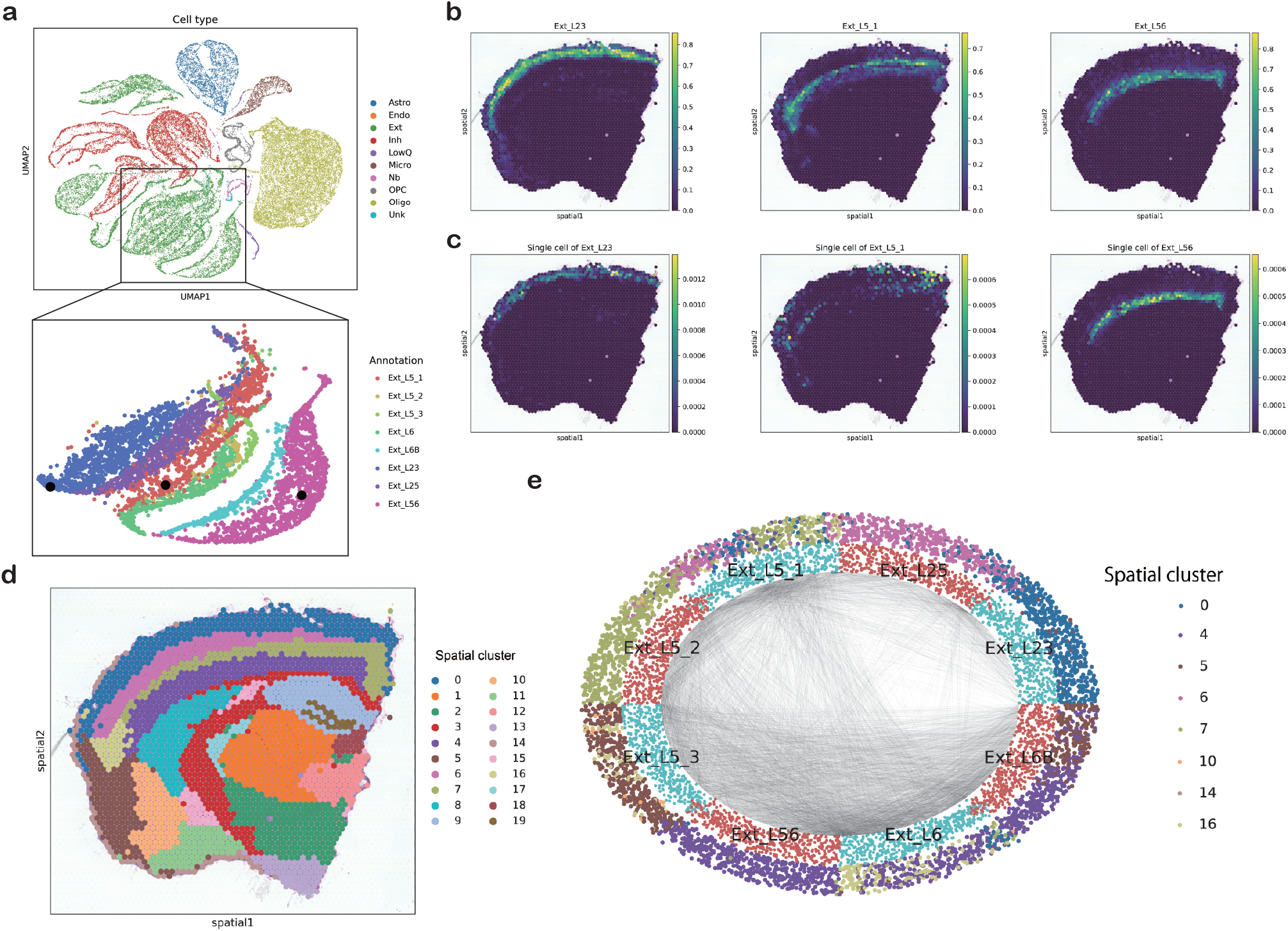
Single-cell decomposition in a mouse cortex dataset. a, UMAP representation of VAE-derived latent states of the single-cell transcriptome. Total (top) and layered excitatory neurons (bottom) are displayed. Black dots in the excitatory neuron panel represent single cells displayed in **c. b**, Spatial assignment of the sub-clusters of three-layered excitatory neurons (Ext L23, Ext L5 1, Ext L56). **c**, Spatial assignment of single cell randomly sampled from three-layered excitatory neuron sub-clusters (Ext L23, Ext L5 1, Ext L56). **d**, Spatial visualization of clustering on the spatial transcriptome. **e**, Visualization of colocalized single-cell pairs (black line) in layered excitatory neurons. Inner layer dots represent a subclass of each single cell. The outer dots represent the most assigned spatial clusters for single cells.

### 3.3 Comparison of accuracy based on the simulated data set

#### 3.3.1 Mapping accuracy for each single cell

We evaluated the accuracy of spatial assignments with DeepCOLOR and other state-of-the-art methods, namely, Cell2location [29], and Tangram [4], based on the simulated dataset of the singlecell and spatial transcriptome. We generated these simulated datasets from a real dataset of squamous cell carcinoma (SCC) [25], assuming five combinations of spatially colocalized cell types. First, we evaluated the assignment amount of truly included single cells to the corresponding spots. We found that DeepCOLOR demonstrated larger assignments on truly included cells than that demonstrated by Cell2location and Tangram (**Fig. 3-a**). For Tangram, the assignment to the majority of truly included cells was close to 0, while there were a few truly included cells with larger assignments than the maximum assignment of DeepCOLOR. This is presumably because Tangram can assign only a few single cells to each spatial spot, and hence estimate distinctively different spatial distribution between single cells with almost the same gene expression profiles. Next, we evaluated the assignment accuracy by the recall of single cells in each spot for specified positive rates and found that the recall values of DeepCOLOR for various positive rates were higher than those of Cell2location and Tangram (**Fig. 3-b**). These results suggested that DeepCOLOR showed superior performance in reconstructing the spatial organization of the single-cell transcriptome without the navigation of cell-type annotation. In contrast, the accuracy of Cell2location was comparable to that of DeepCOLOR when the cell types used for colocalization in the simulation data set were identical to those used for estimation with Cell2location (**Fig. 3-a,b,d,e in Supplementary**). This result indicates that the difference between the reference cell types for estimation and the cell subpopulation in colocalization patterns negatively affected the performance of spatial mapping in cell type-based methodologies. At the same time, DeepCOLOR retained the performance as it was not dependent on the reference cell type information. Next, we validated the ability of DeepCOLOR to extract colocalized cell populations. We calculated the averaged colocalization scores between two cell types and evaluated the accuracy of detecting truly colocalized cell-type pairs using the averaged scores. We found that DeepCOLOR detected the colocalized cell pairs more accurately than the other two methods (**Fig. 3-c**), (**Fig. 3-c,f in Supplementary**). This result suggests that considering the spatial distribution without the cell type label, we extracted colocalized relationships between subpopulations, which is not necessarily consistent with clustering results on scRNA-seq.

**Figure 3:**
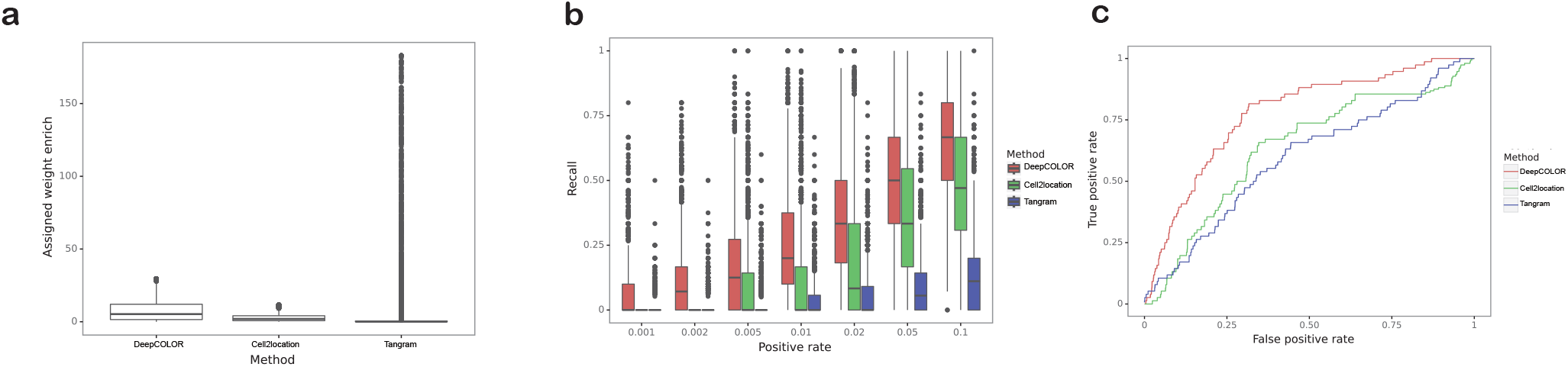
Performance comparison of simulated data. **a**, Enrichment of estimated assignment of single cells originally assigned during simulation. **b**, Recall of originally assigned single cells for a specified positive rate. **c**, ROC curves for detecting cluster pairs belonging to the same region as colocalized population pairs. The resolution parameter of clustering used for the simulation was set to 1.5.

### 3.4 Colocalization network underlying the tumor microenvironment

The tumor microenvironment involves complex CCIs, which significantly affect the prognosis. Here, we applied DeepCOLOR to a dataset of squamous cell carcinoma [25] and explored the colocalization relationships between single cells obtained from the tumor microenvironment. We found that the imputed spatial expression patterns calculated from the transcriptome of the assigned single cells were correlated with the original expression patterns, even for genes that were not used for the assignment estimation (**Fig. 1-b in Supplementary**). We also found that the expression patterns between marker genes for each cell type, CD3D and CD4 for CD4+ T cells and CD8A and GZMB for CD8+ T cells, were more consistent than the raw spatial expression patterns (**Fig. 4 in Supplementary**). Furthermore, the reconstructed spatial expression patterns were smoother than the raw expression counts, while the estimation process did not assume any spatial architecture. These results suggest that DeepCOLOR performed a reasonable single cell assignment for this dataset. The estimated colocalization network among single cells showed that CD1c+ dendritic cells, ASDCs, and Langerhans cells colocalized with heterogeneous tumor keratinocytes referred to as tumor basal keratinocytes (TBKs), tumor differentiating keratinocytes (TDKs), tumor cycling keratinocytes (TCKs), and tumor-specific keratinocytes (TSKs) in the previous study (**Fig. 4-a, b**). All colocalized keratinocyte subpopulations were primarily distributed in spatial transcriptome clusters 7 and 4, the tumor-stromal boundary clusters. In contrast, the TDK and TBK populations distributed in spatial transcriptome cluster 8 demonstrated a low degree of colocalization with any other immune-related cell types and fibroblasts. In contrast, T cells demonstrated strong colocalization with the only subset of TCKs and were distributed only in spatial clusters 2 and 11. These data suggested that several types of dendritic cells, which contribute to the induction of immune responses against tumors, were primarily distributed near the tumor-stromal boundary. In contrast, the T cell population was recruited to the restricted regions in this squamous cell carcinoma. We also noted that the TSK population had colocalized fibroblast populations, which are analyzed in the next section.

**Figure 4:**
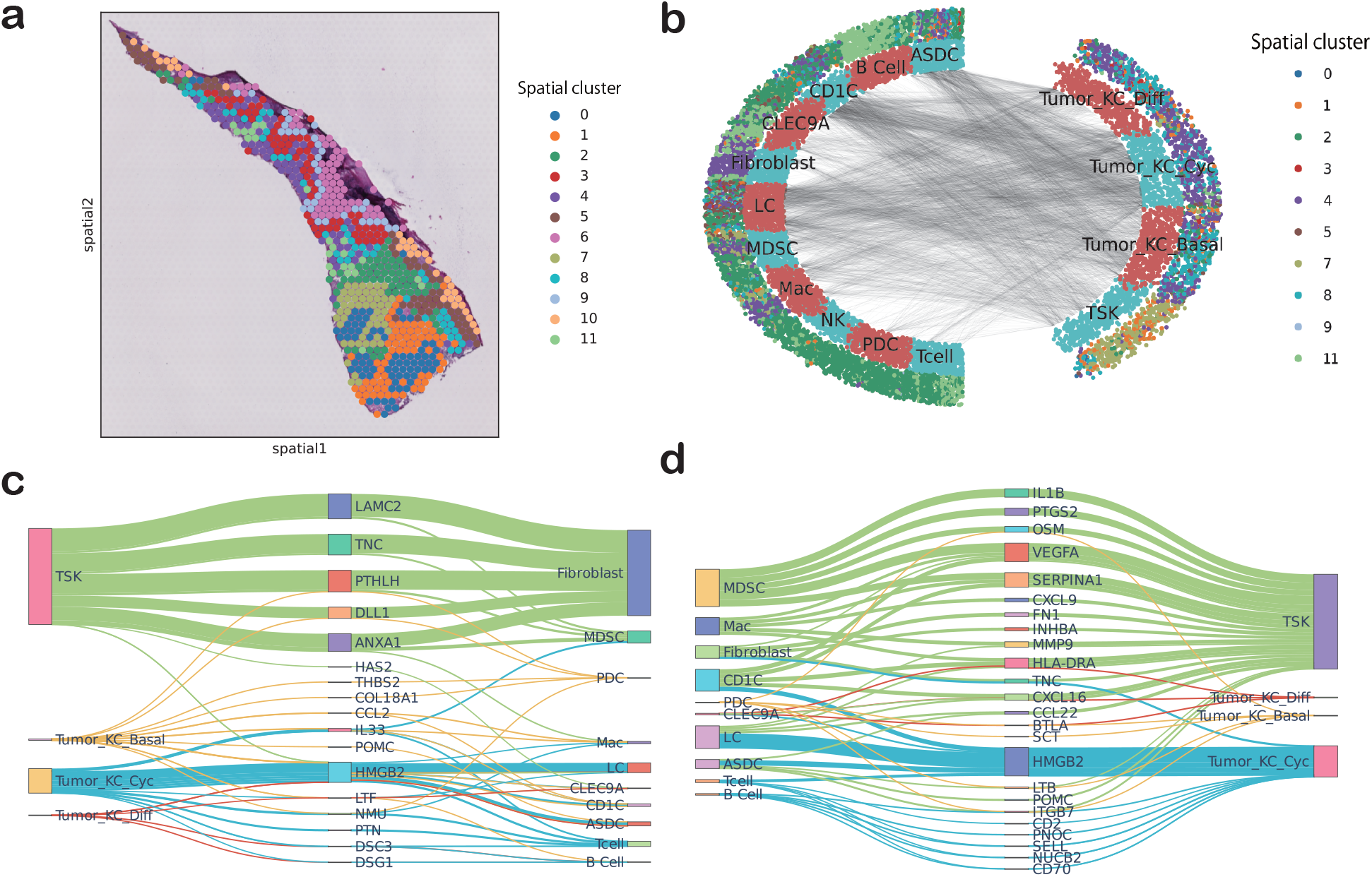
Single-cell decomposition and CCI analysis of an SCC dataset. **a**, Spatial visualization of clustering on the spatial transcriptome. **b**, Visualization of colocalized single-cell pairs (black line) between tumor keratinocytes and immune or stromal cells. Inner layer dots represent the types of single cells. The outer dots represent the most assigned spatial clusters for single cells. **c**, Ligand activity initiating from tumor keratinocytes to immune or stromal cells. **d**, Ligand activity initiating from immune or stromal cells to tumor keratinocytes. The widths of the lines correspond to the ligand activity scores in **c** and **d**. We have only displayed ligands with the top 5 input or output values for each stromal or immune cell type.

Next, we sought to identify the candidates involved in mediating communication between colocalized single cells. Here, we scored ligand activities from tumor subtypes against cells present in the microenvironment (See Methods section) (**Fig. 4-c**). We found that the TNC ligand activity derived from TSKs was the second-highest among the ligand activities against the fibroblast population. TNC expression of cancer cells adjacent to the stroma was reported to correlate with poor prognosis in breast cancer [24], and cultured fibroblasts with TNC treatment was reported to exhibit cancer associated fibroblast like phenotypes [28]. The highest ligand activity against T cell and macrophage populations was that of NMU initiated from TCKs; this was consistent with a previous report showing that NMU secreted by keratinocytes activates various immune cells such as lymphocytes and macrophages and that NMU is associated with tumorigenesis and metastasis [50]. We similarly analyzed the signaling activity initiated from the tumor microenvironment against tumor keratinocytes (**Fig. 4-d**). We found that the CXCL9 activity against TSKs was within the fifth strongest ligand activities initiated from all macrophages, CLEC9A+ DCs, and LCs. CXCL9 is reported to activate invasive and metastatic activities in lung cancer [13]. At the same time, higher CXCL9 expression is associated with tumor depths and positive bone invasion in oral cavity squamous cell carcinoma [8]. In contrast, the activity of INHBA initiated from fibroblasts to TSKs was the second most potent ligand activity observed. This observation is also consistent with a previous report showing that INHBA enhances invasion, proliferation, and growth of gastric cancer cells [10]. Since the candidates of molecular communication machinery observed in our analysis highlighted previously validated molecular communications, these results showed that exploring molecular communications based on single-cell colocalization was an effective approach to dissect the molecular basis underlying the formation of a microenvironment.

#### 3.4.1 Colocalized subpopulations of tumor cells and fibroblasts

Previous analysis of molecular communications based on single-cell colocalization highlighted strong communications between TSKs and fibroblasts. Fibroblasts exhibit various molecular states, which are extensively studied using scRNA-seq technologies [13]. These states exert a significant effect on tumor prognosis [42]. Hence, it is valuable to explore the correlation between these molecular states and heterogeneous cancer cells, some of which demonstrated an invasive leading-edge phenotype in this sample and were termed TSKs in the original analysis. We extracted colocalized populations across fibroblasts and tumor keratinocyte populations including TSKs, based on the single-cell pair colocalization scores (see Methods section). We clustered the colocalized single-cell pairs based on the pair of latent representation and found that tumor cells of the paired cluster 0 had large overlaps with the TSK population (**Fig. 5-a,b**). Furthermore, we confirmed that the spot-wise product of spatial assignments between the colocalized populations were specifically enriched at the tumorstromal boundary (**Fig. 5-c**), which is expected from the leading-edge molecular phenotype of the TSK population. To explore the molecular profiles of the fibroblast population in the paired cluster 0, we analyzed differentially expressed genes in these populations compared to those observed in other fibroblasts (**Fig. 5-d**). These populations demonstrated a high expression of MMP14 associated with processes involved in tumor progression, such as cancer cell invasion via degradation and remodeling of the extracellular matrix [20]. In contrast, INHBA, which was identified as a candidate for molecular communication machinery in a previous analysis, was the second most significantly expressed gene. Here, we also explored the expression patterns of INHBA and its dimer, activin A, in other biological specimens of squamous cell carcinoma by *in situ* hybridization and immunohistochemistry. We found that INHBA expression was enriched in tumor keratinocytes and fibroblasts located at tumor leading edges at both RNA (**Fig. 5-e**) and protein levels (**Fig. 5-f**). Enrichment analysis of positively regulated genes in these populations revealed that the expression of genes involved in glycolysis and hypoxia was upregulated in the TSK-colocalized fibroblast population, pair cluster 0 (*P <* 10^*−*7^ and *P <* 10^*−*10^, respectively). Since the glycolysis pathway is reported to be upregulated in many invasive cancers [19], this result further supports the colocalization of the fibroblast population with TSKs, which demonstrate an invasive leading-edge phenotype.

**Figure 5:**
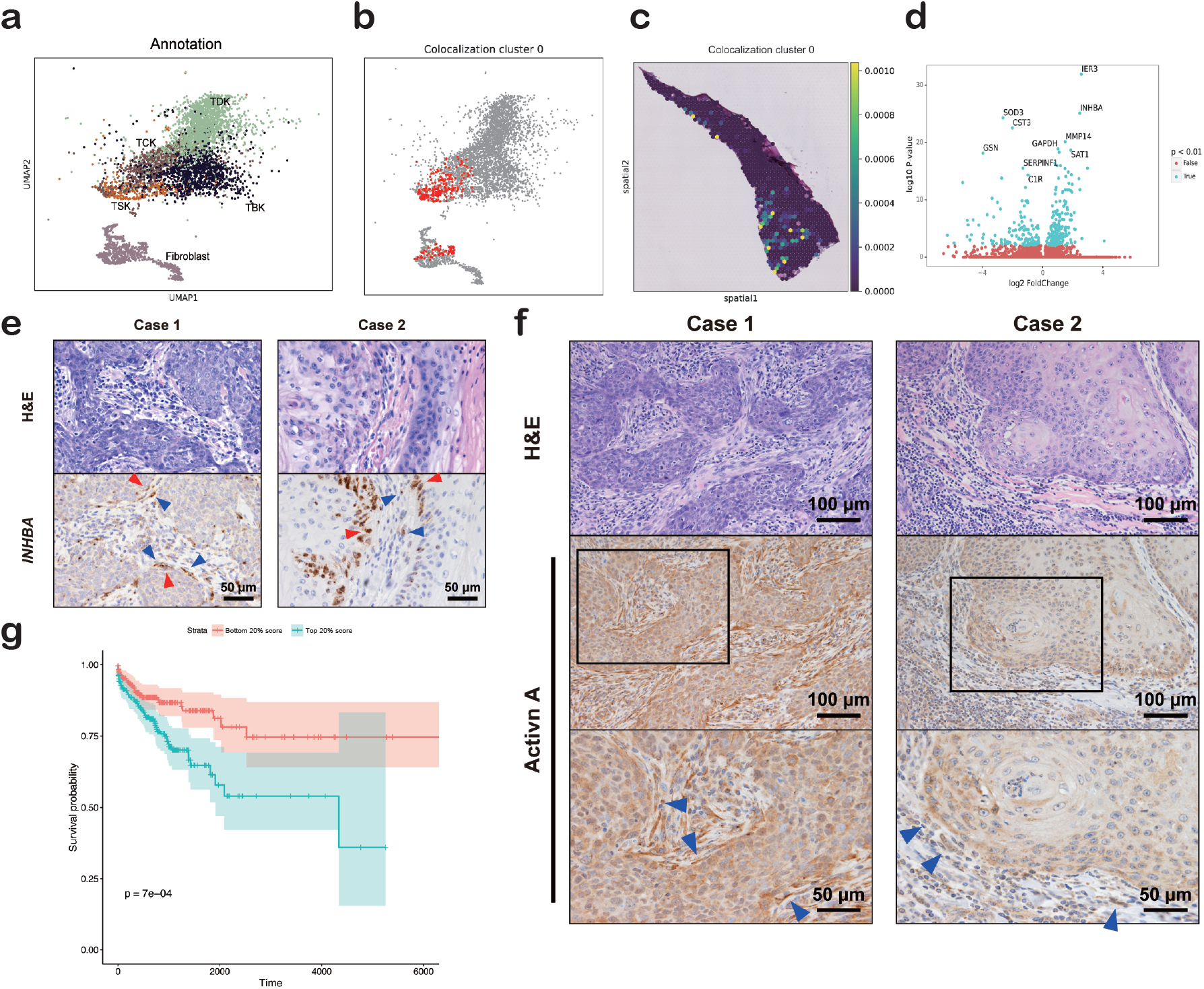
Identification of fibroblast populations colocalized with TSKs. **a**, UMAP representation of VAE-derived latent states of the single-cell transcriptome of tumor keratinocytes and fibroblasts. **b**, Colocalization cluster 0 in UMAP representation. **c**, Spatial distribution of colocalization cluster 0. The spatial distribution was calculated by the summation of assignment products between cells in the colocalization cluster. **d**, Differential gene expression analysis of fibroblasts belonging to colocalization cluster 0 compared to that of the remaining fibroblast population. **e** and **f**, Representative histological images of skin squamous cell carcinoma of two patients. H&E (upper panels in e and f) and ISH with an INHBA-specific antisense probe (lower panels in e) or immunohistochemical staining with anti-activin A antibody (middle and lower panels in f) in the same area in serial sections. Boxed areas in the middle panels in e are presented as magnified images in the lower panels. INHBA-positive cancer cells are indicated by red arrowheads and INHBA-positive or activin A-positive cancer-associated fibroblasts are indicated by blue arrowheads. **g**, Kaplan-Meier plot of the survival rate for patients with SCC with high and low signature scores of fibroblasts belonging to colocalization cluster 0 (top and bottom 20%) in TCGA dataset.

#### 3.4.2 Reproducibility analysis of ST and TCGA

We investigated whether these patterns can be analyzed via spatial gene expression patterns derived from other observation technologies to validate the colocalization patterns between cancer cells and fibroblasts. In particular, we decomposed the spatial gene expression patterns observed in the same study by spatial transcriptomics (ST) into the same single-cell gene expression profiles obtained using DeepCOLOR. We extracted cancer cells which form colocalized pairs in ST deconvolution with fibroblasts that belong to the previously identified colocalization cluster 0 in Visium deconvolution. The recovered ST-colocalized tumor cells significantly overlapped with tumor cells that belonged to the colocalization cluster 0 (odds ratio 6.70). These results indicated that DeepCOLOR was able to reproducibly identify unique colocalization patterns between tumor cells and fibroblasts identified in Visium via another spatial transcriptome observation technique, ST. Next, we investigated whether this colocalized population is likely to co-occur across many patients and associated with the prognosis difference. The signature scores for the colocalized tumor cells and fibroblasts in the colocalization cluster 0 were positively correlated (Pearson’s correlation was 0.765) across the transcriptome of patients with SCC derived from TCGA (**Fig. 5 in Supplementary**). Furthermore, the signature scores of colocalized fibroblasts exhibited a strong association with worse overall survival (*P* = 0.0007) (**Fig. 5-g**). However, the association between the signature scores of the colocalized tumor cell population and worse overall survival was relatively moderate (*P* = 0.21) (**Fig. 6 in Supplementary**). These findings indicate the identified colocalized populations between tumor cells and fibroblasts exist in various patients with SCC. Furthermore, the stronger association of the colocalized fibroblast population with prognosis indicates that the arrival of fibroblasts to this colocalization niche enhances the malignancy of the tumor.

**Figure 6:**
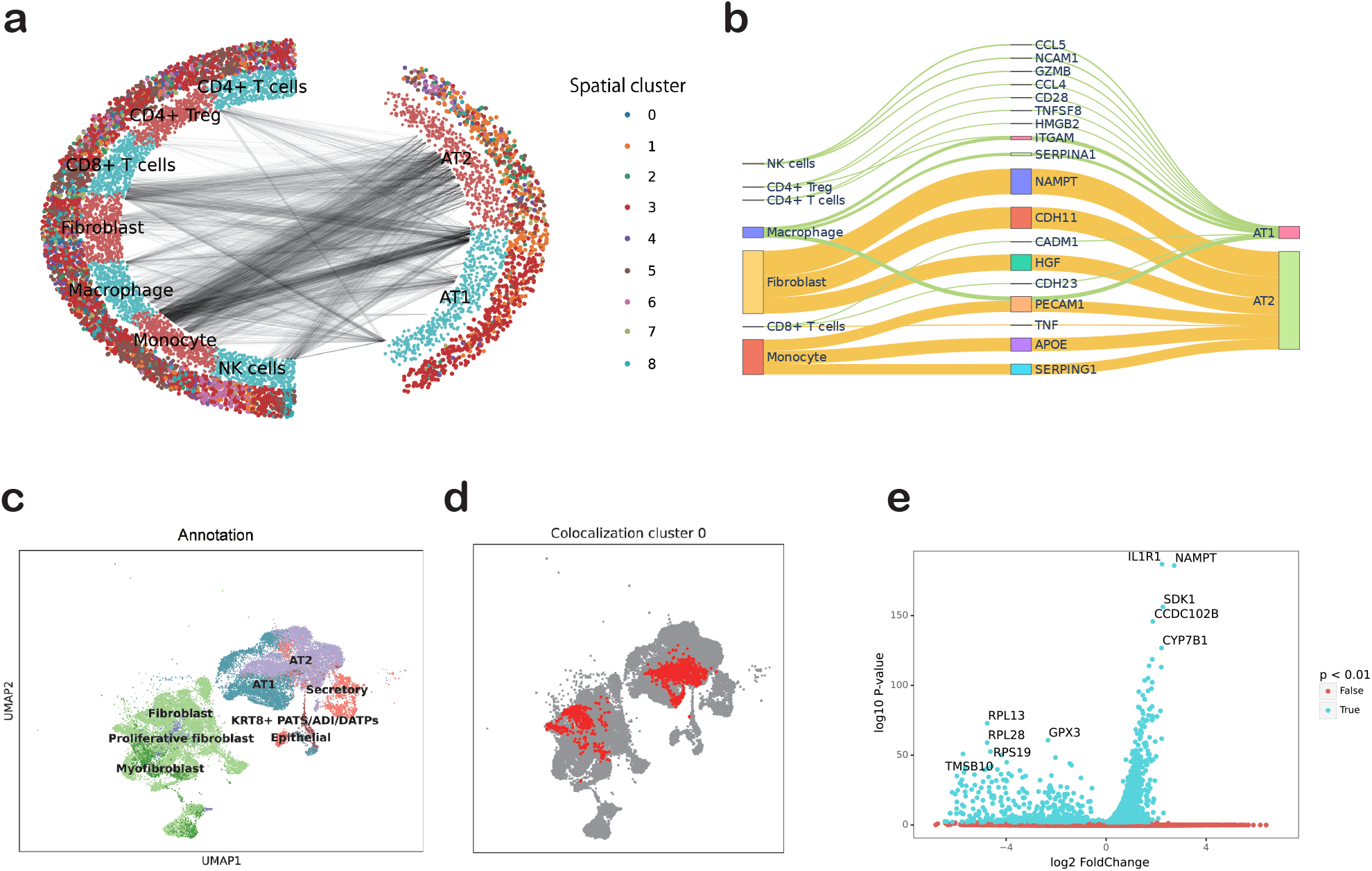
Single-cell decomposition and CCI analysis of a COVID-19 dataset. **a**, Visualization of colocalized single-cell pairs (black line) between alveolar cells and immune or stromal cells. Inner layer dots represent the types of single cells. The outer dots represent the most assigned spatial clusters for single cells. **b**, Ligand activity initiating from immune or stromal cells to alveolar cells. The widths of the lines correspond to the ligand activity scores. We have only displayed ligands with the top 5 input or output values for each stromal or immune cell type in **b. c**, UMAP representation of VAE-derived latent states of the single-cell transcriptome of epithelial cells and fibroblasts. **d**, Colocalization cluster 0 in UMAP representation. **e**, Differential gene expression analysis of fibroblasts belonging to colocalization cluster 0 compared to that of the rest of fibroblast populations.

### 3.5 SARS-CoV-2

#### 3.5.1 Alveolar type II cells colocalized with macrophages in SARS-CoV-2 infection

We next applied DeepCOLOR to another dataset of SARS-CoV-2 [12] composed of single-cell transcriptome observations and spot-wise transcriptome observations in lung tissues of SARS-CoV-2 patients. We estimated the quantitative assignment of single cells to the spots and reconstructed gene expression of the spots based on that of single cells assigned to the spots. The reconstructed expression patterns for the spots were well correlated with true expression patterns, even for genes that were not used for the estimation (**Fig. 1-c in Supplementary**). When we visualized the colocalization between alveolar cells and surrounding cells, such as immune cells and fibroblasts, we found that a specific subpopulation of alveolar type II (AT2) cells, the abundance of which is primarily reduced in patients with severe COVID-19, demonstrated a remarkable colocalization with various cell types; this population was mainly distributed to spatial cluster 1 (**Fig. 6-a**). The spots belonging to spatial cluster 1 were mainly annotated as PanCK+ alveolar (38 of 59), which were associated with SARS-CoV-2 infection [23]. To quantify the molecular communication between the surrounding cells and alveolar cells, we calculated the ligand activity between surrounding cells and alveolar cells based on the estimated single-cell colocalization (**Fig. 6-b**). We found that the strongest ligand activity initiating from fibroblasts to AT2 cells was that of NAMPT, which plays an important role in the activation of the innate immune response [7] and is associated with the development of acute respiratory distress syndrome in lung injury [40]. The activity of PECAM1 was the strongest activity initiating from monocytes to AT2 cells, while the expression level of PECAM1 was associated with the severity of COVID-19 [31]. We also found that the strongest ligand activity initiating from CD8+ T cells to AT2 cells was that of TNF, the expression level of which is also associated with disease severity and survival of patients with COVID-19 [11]. These results showed that the colocalization-based ligand activity analysis discerned appropriate candidates for molecular communication among patients with COVID-19. Next, we analyzed singlecell colocalization between epithelial cells and fibroblasts, dissecting the most potent ligand activity toward AT2. We extracted one of the colocalization clusters (cluster 0) with large overlap with AT2 cells (**Fig. 6-c,d**). We analyzed the gene expression profiles of the fibroblast population in the colocalization cluster and found that NAMPT, which demonstrated a strong ligand activity to AT2 cells, was the second most significantly enriched gene in the colocalized fibroblast cluster (**Fig. 6-e**). We also found that the pathway activity of oncostatin M, the expression of which was reported to be elevated in the serum of patients with COVID-19 [35], was significantly enriched (*P <* 10^*−*4^). At the same time, the most significantly upregulated gene was IL1R, the ligand of which is an important marker of severe symptoms among patients with COVID-19 [11]. These data suggest that the fibroblasts received molecular signals responsible for severe symptoms and acquired a molecular phenotype, contributing to the severity in patients with COVID-19.

## 4 Discussion

This article presents a new deep learning framework called DeepCOLOR, which enabled us to analyze colocalization networks across single cells with deep molecular profiles captured by scRNA-seq. This new computational framework showed higher accuracy for mapping scRNA-seq observation to spot-level spatial transcriptome data and detecting colocalized cell populations than existing methods in simulation experiments and demonstrated a finer anatomical distribution than cell-type distribution due to its label-free approach. Furthermore, DeepCOLOR extracted plausible candidates involved in the molecular machinery underlying cell-cell communication and disease-associated colocalized populations in a squamous cell carcinoma dataset [25] and COVID-19 dataset [12]. In particular, DeepCOLOR highlighted the molecular communication machinery consistent with the disease phenotype in both datasets. Our analysis predicted that the expression of INHBA, associated with enhanced invasion, was enriched in fibroblasts colocalized with invasive tumor cells. This prediction was validated by detecting both protein and RNA expression in biological samples independent from the dataset used for the estimation. These results highlight the significance of single-cell-level colocalization relationships for dissecting molecular communications underlying disease progression.

Recently, spot-level spatial transcriptome observation is garnering significant interest and is being used for various biological systems, including the tumor microenvironment [32, 39]. While these observations are useful for associating visualized tissue phenotypes with molecular phenotypes, it would be difficult to dissect a complex molecular process mediated across various cells. For this purpose, researchers developed many computational methodologies integrating the spatial transcriptome with corresponding scRNA-seq observations [1, 4, 6, 16, 29]. However, most of these methodologies relied on cell type labeling with scRNA-seq observations, which can be the upper resolution limit for spatial distribution analysis. Indeed, we showed that single cells could be spatially assigned to more specific regions compared with cell type mapping of the fine grained excitatory neuron populations. Furthermore, simulation experiments for the integration of single cell and spatial transcriptome showed that, for cell type based spatial deconvolution, the deviation between the true population structures and the assumed population structures induces an accuracy decay for both spatial assignment of single cells and prioritization of colocalization relationships. Hence, the label-free property of DeepCOLOR enabled us to capture the colocalization network of various cell populations, identifying niche environments with unprecedented accuracy.

Cell-cell communications are crucial for not only normal development but also disease progression [3]. Indeed, many recent advancements in therapeutic strategies involve the perturbation of cellcell communication [47]. Hence, computational methodologies inferring cell-cell communication are extensively developed by targeting scRNA-seq observations [5,15,26]. However, single-cell molecular profiles derived from scRNA-seq observation lose their spatial context. Adequate spatial proximity is an important factor contributing to the induction of cell-cell communication. We quantified ligand activity only between colocalized single-cell pairs, which are more likely to communicate with each other than randomly selected single-cell pairs. This analysis highlighted several pathways of cell-cell communication reported in previous studies and predicted novel molecular machinery involved in mediating communication between fibroblasts and invasive tumor cells, which was validated at the protein and RNA level.

A major limitation of DeepCOLOR is that it deconvolutes all spatial transcriptome profiles into single-cell molecular profiles captured by scRNA-seq. Hence, if spatial transcriptome-specific cell populations do not exist in scRNA-seq, the deconvolution of spatial transcriptome spots, including the populations, would be unreliable. One possible solution for this issue would be assuming several pseudo inputs for scRNA-seq, which is optimizable and expected to be similar to the molecular profiles of the populations unobserved in scRNA-seq since such pseudo-inputs approach succeeded in capturing population structure as prior means in a variational auto encoder with VampPrior [45]. The improvements on this issue would increase the reliability of DeepCOLOR for the unpaired spatial transcriptome and single-cell transcriptome, even if the included cell populations of them could have some discrepancy.

Recent advancements in the observation of the single-cell transcriptome with other modalities, such as open chromatin and the proteome, are opening new avenues to analyze the cross-talk between the different layers of biological processes at the omics scale [9, 34]. For cell-cell communication, protein signals from other cells alter the epigenetic profiles of the nucleus and change the transcription kinetics of various genes, which generate molecular signals for communicating with other cells through the protein layer. Hence, the application of DeepCOLOR to such multimodal single omics data with spatial transcriptome is expected to produce a concrete basis for detecting molecular changes in various layers induced by CCI. Finally, we anticipate that our newly developed computational framework could be utilized for uncovering molecular pathways via different molecular layers and single-cell colocalization networks.

## 5 Methods

### 5.1 Data preprocessing and downstream analysis

Using ‘scanpy’ Python package [48], we excluded single cells and spatial spots which expressed fewer genes than 500 genes or more mitochondrial genes than 5% of total expression. We conducted and visualized UMAP embeddings of the latent cell states of single cells using ‘scanpy’. We also utilized ‘scanpy’ for clustering spatial transcriptome data by the Leiden clustering algorithm with default parameters.

### 5.2 Spatial mapping of single cells using a neural network

To quantify the expected contribution of every single cell in scRNA-seq for determining all spatial spots in the spatial transcriptome, we employed a probabilistic model for the spatial transcriptome observation, given the expected contribution of all single cells observed in the scRNA-seq analysis. We estimated the contribution by maximizing the likelihood of the probabilistic model for observation of the spatial transcriptome. However, the naive formulation of this problem can lead to overfitting due to the numerous independent parameters. We employed a continuous mapping function from a latent representation of a single cell to a spatial spot to overcome this limitation. This formulation imposed a constraint on the mapping in that single cells with almost the same molecular profiles were mapped similarly. This constraint was enhanced by the stochastic gradient descent, where down-sampled single cells were used to calculate the likelihood gradient. This section introduces a variational autoencoder (VAE) for obtaining a stochastic latent representation of the single cells, the probabilistic model of the spatial transcriptome, and the optimization procedure.

#### 5.2.1 Derivation of the stochastic latent representation of the single cell transcriptome

We utilized a variational auto encoder (VAE) for deriving latent representations for single cell transcriptome observations. We defined the generative model of scRNA-seq observation of the cell *c, x*_*c*_ *∈* ℝ^*G*^ as shown below:

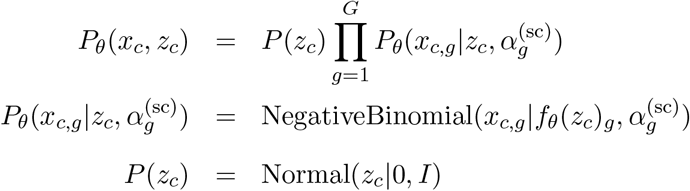

where *G* is the number of genes, *z ∈* ℝ^*M*^ is a latent cell state and *f*_*θ*_ : ℝ^*M*^ *→* ℝ^*G*^ is a decoder neural network described in **Supplementary Table 1** and 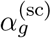 is the dispersion parameter of the gene *g*. We approximated the posterior distribution of latent representation *P* (*z*_*c*_|*x*_*c*_) *∝ P* (*x*_*c*_, *z*_*c*_) using the Gaussian distribution as shown below:

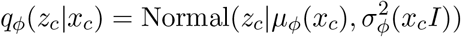

where 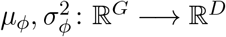 are encoder neural networks described in **Supplementary Table 1**. To approximate the true posterior distribution appropriately, we maximized the evidence lower bound (ELBO) for *θ* and *φ*, which is defined as follows:

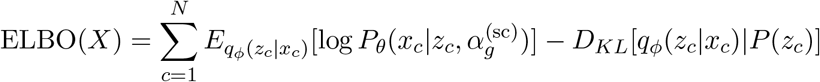

where *X* = (*x*_1_, …, *x*_*N*_) ^*T*^ and *N* is the total number of cells. We maximized this ELBO using the Adam optimizer implemented with a learning rate of 0.0004 for 500 epochs.

#### 5.2.2 Probabilistic model of spatial transcriptome data

We assumed that the expression of gene *g* at spatial spot *s, e*_*s,g*_ follows a negative binomial distribution as shown below:

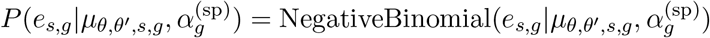

where 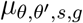 is the unobserved expression level of the gene *g* at spot *s* and 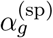 is the dispersion parameter of the gene *g*. To attribute the spatial expression profile to the expression profile observed using scRNA-seq, we constructed an expected contribution of cell state *z* for spot *s* as a continuous function implemented by neural network 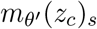 described in **Supplementary Table 1**. Using the mapping function, we modeled 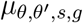 as the weighted average of the scRNA-seq expression profile, given the following approximated posterior distribution of the latent cell states:

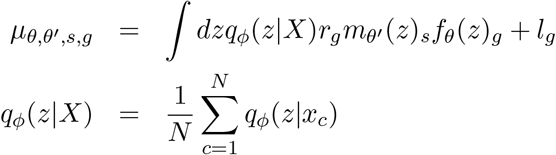

where *q*_*φ*_(*z*|*X*) is the posterior distribution of a latent cell state, given the total scRNA-seq data set *X, r*_*g*_ is the gene-wise technical capturing ratio of spatial transcriptome observation compared to that of scRNA-seq, and *l*_*g*_ is the gene-wise shift parameter that is assumed to represent ambient RNA in the spatial transcriptome data. Since the exact integration in equation X is not feasible, we calculated the stratified Monte Carlo approximation of the posterior distribution as shown below:

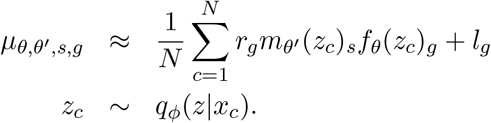

#### 5.2.3 Stochastic optimization for smooth mapping function

To derive the mapping function optimized for the data, we maximized the log likelihood

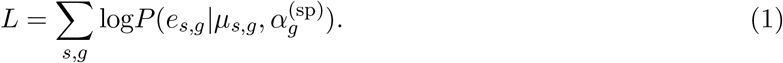

Since the computational complexity of the mean parameter defined above is proportional to the number of cells, we calculated and optimized the likelihood for spatial transcriptome observation with the mean parameter for mini-batches of single cells, *M* :

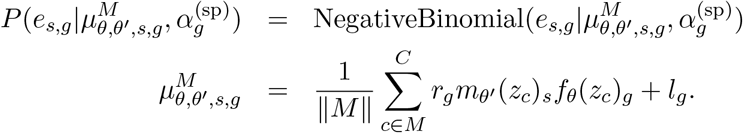

This downsampling for single cells imposed a constraint on the mapping function in that the random subsets of single cells could reconstruct consistent expression profiles of spatial observations. Hence, the mapping function was expected to be enhanced for estimating similar mapping profiles for single cells with similar latent representation. To optimize this stochastic likelihood, we utilized Adam implemented in PyTorch with a learning rate of 0.0004 for 500 epochs. For this optimization, we did not used randomly selected 2% of cells and 10% of genes for testing the accuracy. We also note that we did not update the parameters of *θ* and *φ* so that the encoder and decoder networks keep the information on single-cell expression profiles.

### 5.3 Colocalization analysis based on spatial mapping

#### 5.3.1 Construction of the expected colocalization matrix

DeepCOLOR could estimate the contribution of every single cell to each spot 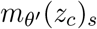. We utilized this property for the analysis of colocalization among single cells. First, to filter out single cells that were not mapped well, we excluded cells whose cumulative values of total contribution 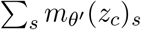 were lower than 0.05. Next, we normalized the spatial distribution for each cell so that the summation for all spots was equal to 1, 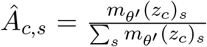. We calculated the colocalization matrix as a product of the normalized spatial distribution and its transpose:

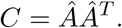

Here, the element of the colocalization at *i*th row and *j*th column represents the colocalization score between cell *i* and cell *j*. We calculated the log ratio of the scores to that observed between two cells that were uniformly mapped to each spatial spot:

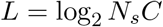

where *N*_*s*_ is the number of spots in the spatial transcriptome. We selected all colocalization pairs *i, j* whose *L*_*i,j*_ exceeded 1. This criterion for the colocalization pair corresponds to the case where pairs were localized together with a probability of two times higher than that observed when uniformly distributed across all spots.

#### 5.3.2 Ligand activity between colocalized single cells

To dissect the molecular machinery involved in mediating cell-cell communication, we combined the ligand-target regulatory potential implemented within Nichenet with the expression profiles of colocalized cell pairs [5], representing how strongly existing knowledge supports the influence of the ligand on the expression of the target gene in other cells, with the detection of ligand expression of colocalized cells. We calculated the receiver scores for ligands *l* as follows:

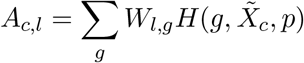

where *W*_*l,g*_ is a ligand–target regulatory potential value of Nichenet [5], *H*(*i, x, p*) = *I*(*x*_*i*_ *> q*_*p*_(*x*)), *q*_*p*_(*x*) is the *p* quantile of vector *x* and 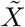 is the scaled expression that was calculated using the ‘scanpy.pp.scale’ function implemented in the Scanpy package after resampling 500 cells per every cell cluster to be analyzed [48]. We calculated the colocalized ligand activity of *l* from single cell cluster *k* to *k*^*′*^:

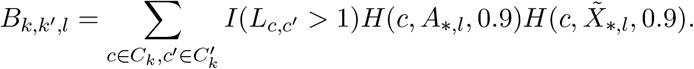

where *C*_*k*_ is *R* resampled single cells of cluster *k* (*R* = 500 in this study) and 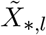 and *A*_*∗,l*_ denotes *l*-th column vector of *X* and *A*. The colocalized ligand activity 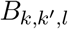 corresponds to the expected number of colocalized cell pairs with high ligand expression and high ligand activity between the cell types.

#### 5.3.3 Clustering colocalized pairs of single cells

We derived the latent representation of colocalized pairs described above as the summation of the single-cell latent representations, which were derived from a VAE of scRNA-seq 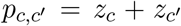. We extracted the colocalized pairs between two clusters of single cells subjected to colocalization analysis. We clustered the latent representation of colocalized pairs using the Leiden algorithm implemented in Scanpy with resolution parameter 0.1. This clustering of the latent representations of colocalized pairs segregated the subpopulation of colocalized pairs with similar molecular profiles.

#### 5.3.4 Differentially expressed gene analysis of colocalization clusters

To characterize the molecular profiles of single cells that belonged to specific colocalization clusters, we conducted a differentially expressed gene (DEG) analysis between single cells in the colocalization clusters and the other single cells that belonged to the same single cell clusters. We used the Wilcoxon rank-sum test with Benjamini-Hochberg multiple test correction for this DEG analysis, implemented in the Scanpy dataset. For gene enrichment analysis, we conducted Fisher’s exact test for gene sets recorded in IMPaLA [27].

#### 5.3.5 TCGA analysis of the correlation and the survival date association of colocalized population signatures

To determine whether the identified colocalized population was also colocalized across samples, we analyzed the correlation between the signature scores of both populations in the TCGA dataset. We used FPKM values of RNA-seq data obtained in 1993 from squamous cell carcinoma samples to calculate the population signature scores. The signature score for each sample was calculated using the mean z-scores of log FPKM values plus 1 of population-specific genes. The population-specific genes with up-or down-regulated expression were defined as genes with adjusted p-values smaller than 0.01 and log2 fold changes larger than 1 or smaller than −1, as observed in DEG analysis for each population. We excluded overlapped genes for the calculation of signature scores. We evaluated Pearson’s correlation between the signature scores for both populations. The association between survival date and these scores and gene expression levels was analyzed using R package survival after stratification into the top 20% and bottom 20% of scores and expression levels.

### 5.4 Simulation of the spatial transcriptome

To evaluate the performance of DeepCOLOR in the spatial assignment of the single-cell transcriptome and detection of colocalized populations, we conducted a simulation of the spatial transcriptome from reference scRNA-seq data similar to the simulation method implemented in [29]. First, we separated the scRNA-seq population into two randomly selected subpopulations for simulation and training, defined as *C*^(*s*)^ and *C*^(*t*)^, respectively. Next, we assumed *R* = 10 regions, each composed of randomly selected clusters from *K* clusters of scRNA-seq data derived by ‘scanpy.tl.leiden’ with specified resolution parameters. We determined the abundance of region *r* in spot *s* as 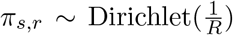. The composition of cluster *k* in region *r* was 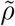. where 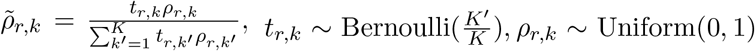 and *K*^*′*^ = 5. Combining these two hierarchical compositions, we calculated the expected abundance of single cells *c* of the simulation dataset in each spot *s* as follows:

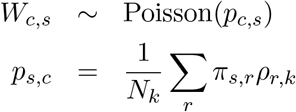

where *c* belongs to cluster *k*, and *N*_*k*_ represents the number of cells in cluster *k* used for the simulation dataset. We simulated the spatial gene expression of gene *g* at spot *s* from the weighted average of single cell expression profile *X*:

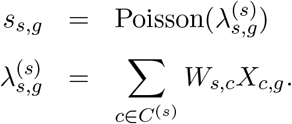

For evaluation, we assigned the weight values of cell *c* in the simulation dataset to the nearest neighbor cell in the training dataset *N* ^(*s,t*)^(*c*), based on 30 dimensional PCA-coordinates of their expression profiles. Hence, the assignment of cell *c*^*′*^ of the training population in spot *s* is

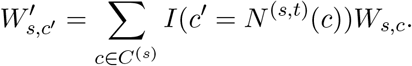

#### 5.4.1 Evaluating spatial assignment of single cells

For the evaluation of spatial assignment, we calculated the recall of the training cells that were most similar to simulation cells included in each spot

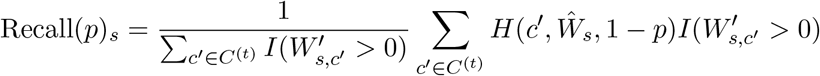

where *p* is the specified positive rate and 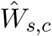 is the estimated assignment of cells *c* in spot *s*.

#### 5.4.2 Evaluating detection of colocalized cell populations

For the detection of colocalized populations, we evaluated the detection accuracy of cluster pairs belonging to the same region. As a predictor, we calculated the mean colocalization scores across cell pairs within each cluster pair:

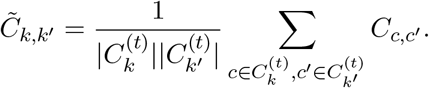

#### 5.4.3 Comparison with other methods

We compared the performance of DeepCOLOR with that of existing computational methodologies for deconvolving spot-wise spatial transcriptomes, namely, Cell2location [29], and Tangram [4]. For both methodologies, we used default parameters used in the evaluation experiments. Since Cell2location provides cluster-wise abundance for each spot, we deconvolved the weights into every single cell equally for performance evaluation.

### 5.5 Human skin squamous cell carcinoma samples and histological analysis

Surgically resected skin samples from patients with squamous cell carcinoma were obtained at Nagoya University Hospital. This study was approved by the Ethics Committee of Nagoya University, Graduate School of Medicine. Human tissues were fixed in 10 % neutral-buffered formalin, dehydrated, and embedded in paraffin. Immunohistochemical analysis was performed using anti-Activin A antibody (Novus Biologicals, Centennial, CO, USA) as previously described [38]. In situ hybridization (ISH) analysis was performed by RNAscope technology (RNAscope 2.5 HD Detection Kit; Advanced Cell Diagnostics, Newark, CA, USA). Briefly, human tissue sections were baked in an oven at 60°C for 1 h, deparaffinized, and incubated with H2O2 solution for 10 min at room temperature. The slides were boiled in target-retrieval solution for 3 min in a pressure cooker (SR-MP300; Panasonic, Kadoma, Japan) and incubated with protease solution for 30 min at 40°C. The slides were then incubated with the relevant probe (human INHBA, NM 002192.4, region 337-3141; Advanced Cell Diagnostics) for 3 h at 40°C in a dry oven (HybEZ II Hybridization System; Advanced Cell Diagnostics), followed by successive incubation with Amp1-6 reagents. The staining was visualized with 3,3’-diaminobenzidine, followed by counterstaining with hematoxylin. Two independent pathologists evaluated the human tissues subjected to ISH and hematoxylin and eosin (H&E) staining.

### 5.6 Data and code availability

We derived combined spatial and single-cell transcriptome datasets from Gene Expression Omnibus (Mouse brain cortex dataset:, SCC dataset: GSE144240, and SARS-CoV2 dataset: GSE171668).

Codes for our analysis, including DeepCOLOR, are available at https://www.github.com/kojikoji/deepcolor.

## 6 Acknowledgement

This was supported by The Japan Society for the Promotion of Science (grant no. 20H04841, 20H04281, and 20K21832), The Japan Agency for Medical Research and Development (AMED) (grant no. JP20ek0109488 and JP21wm0425007) and The Japan Science and Technology Agency (JST) (Moonshot R&D, grant no. JPMJMS2025). The computational resource of AI Bridging Cloud Infrastructure (ABCI) was provided by the National Institute of Advanced Industrial Science and Technology (AIST), and SHIROKANE by the Human Genome Center, the University of Tokyo.

## 7 Author contributions

YK designed the study and performed software development and data analysis. SM and AE conducted and supevised experiments. SH, HH and MI conducted software development and data analysis. AE collected samples. TS designed and supervised the study.

