## Supplementary figures for "Single-cell colocalization analysis using a deep generative model"

Table 1: deepCOLOR architecture

| Name | Operation | Dimension | Layer Normalization | Activation | Input |
|---|---|---|---|---|---|
| **Inputs** | | | | | |
| SCT | - | #Genes | - | - | - |
| **Encoder** | | | | | |
| $h_1^{(e)}$ | FC | 50 | Yes | ReLU | SCT |
| $h_2^{(e)}$ | FC | 50 | Yes | ReLU | $h_1^{(e)}$ |
| $\mu(x)$ | FC | 10 | No | - | $h_2^{(e)}$ |
| $\sigma(x)$ | FC | 10 | No | Softplus | $h_2^{(e)}$ |
| $z$ | Sample from Normal | 10 | - | - | $\mu(x)$, $\sigma(x)$ |
| **Decoder** | | | | | |
| $h_1^{(d)}$ | FC | 50 | Yes | ReLU | z |
| $h_2^{(d)}$ | FC | 50 | Yes | ReLU | $h_1^{(d)}$ |
| $\lambda(x)$ | FC | #Genes | No | Softplus | $h_2^{(d)}$ |
| **Distributer** | | | | | |
| $h_1^{(s)}$ | FC | 50 | Yes | ReLU | z |
| $h_2^{(s)}$ | FC | 50 | Yes | ReLU | $h_1^{(s)}$ |
| $m(z)$ | FC | #Spots | No | Softplus | $h_2^{(s)}$ |

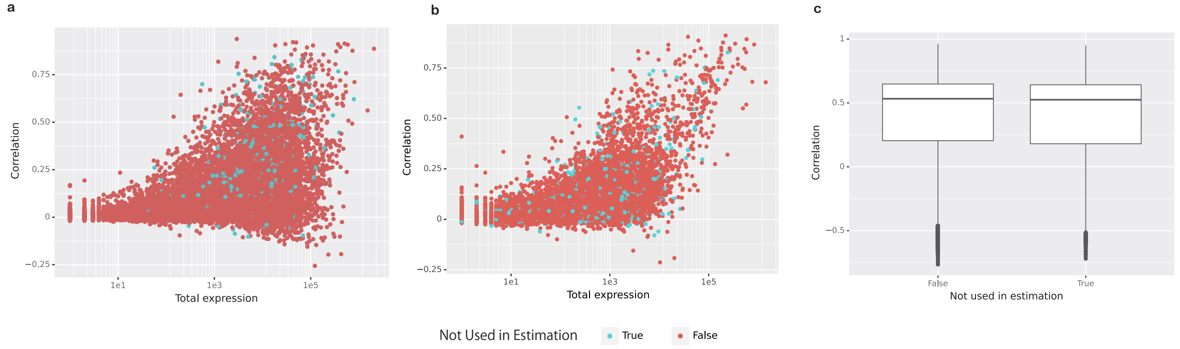


Figure 1: **Correlation between original and reconstructed spatial gene expression in mouse cortex dataset.** We calculated gene-wise Pearson's correlation. Reconstructed spatial gene expression is calculated by the weighted average of single cell expression decoded from VAE, based on the estimated spatial assignment. We displayed the correlation scores with the total expression level of each gene for mouse cortex dataset, **a**, SCC data set, **b**. We displayed the correlation score distribution for Covid19 dataset, **c**.

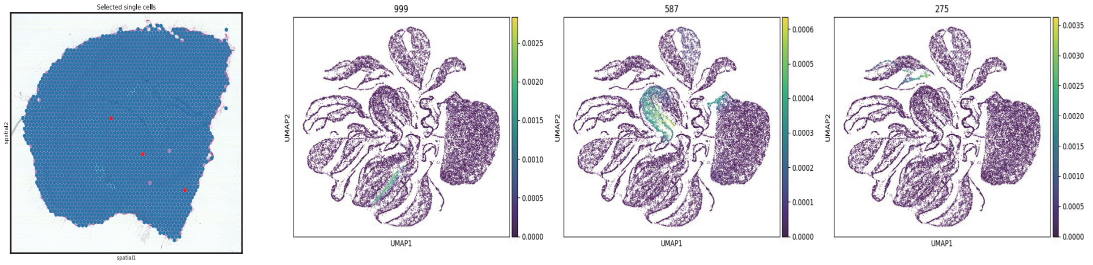


Figure 2: **Assignments of single cells for randomly selected spatial spots.** Randomly selected spatial spots are displayed in the most left panel. Three right panels displays assignment for the three spatial spots.

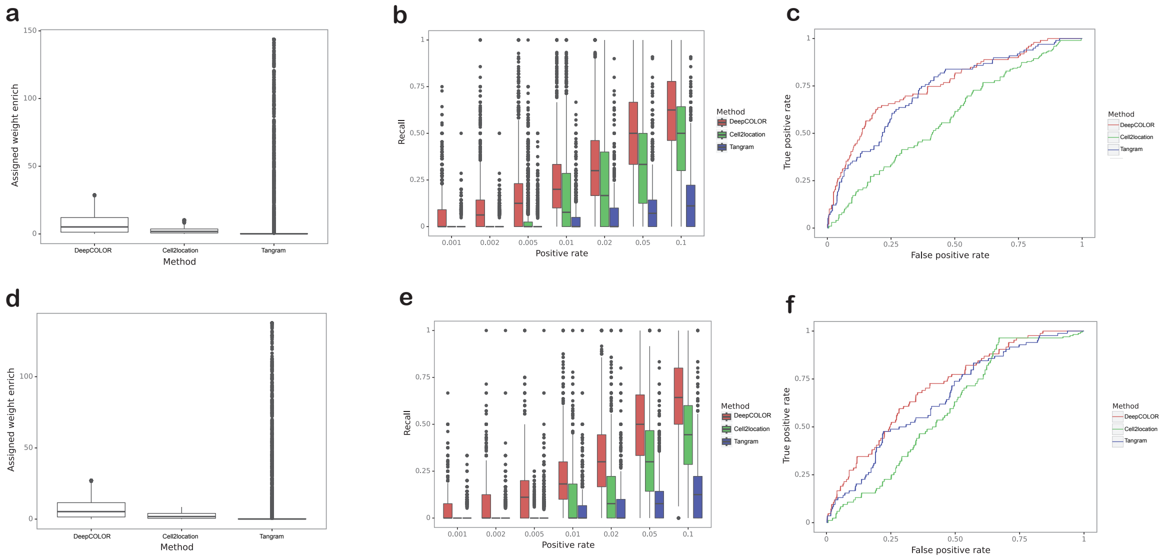


Figure 3: **Performance comparison on simulated data** a, d, Enrichment of estimated assignment to single cells originally assigned in simulation. b, e, Recall of originally assigned single cell for specified positive rate. c, f, ROC curves for detecting cluster pairs belongs to a same region as colocalized population pairs. Resolution parameter of clustering used in simulation was set to 1.0 in a, b and c. Resolution parameter of clustering used in simulation was set to 0.5 in d, e and f.

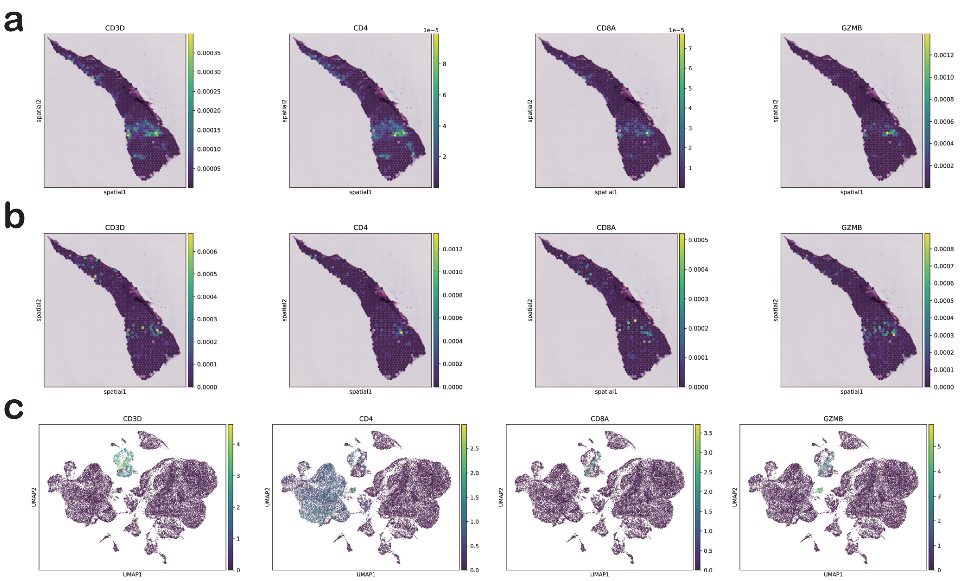


Figure 4: **Raw expression and reconstructed expression of marker genes** a, Reconstructed spatial expression patterns for CD3D, CD4, CD8A and GZMB. b, Raw spatial expression patterns for CD3D, CD4, CD8A and GZMB. c, Single cell expression patterns for CD3D, CD4, CD8A and GZMB in UMAP representation of cell states derived from VAE.

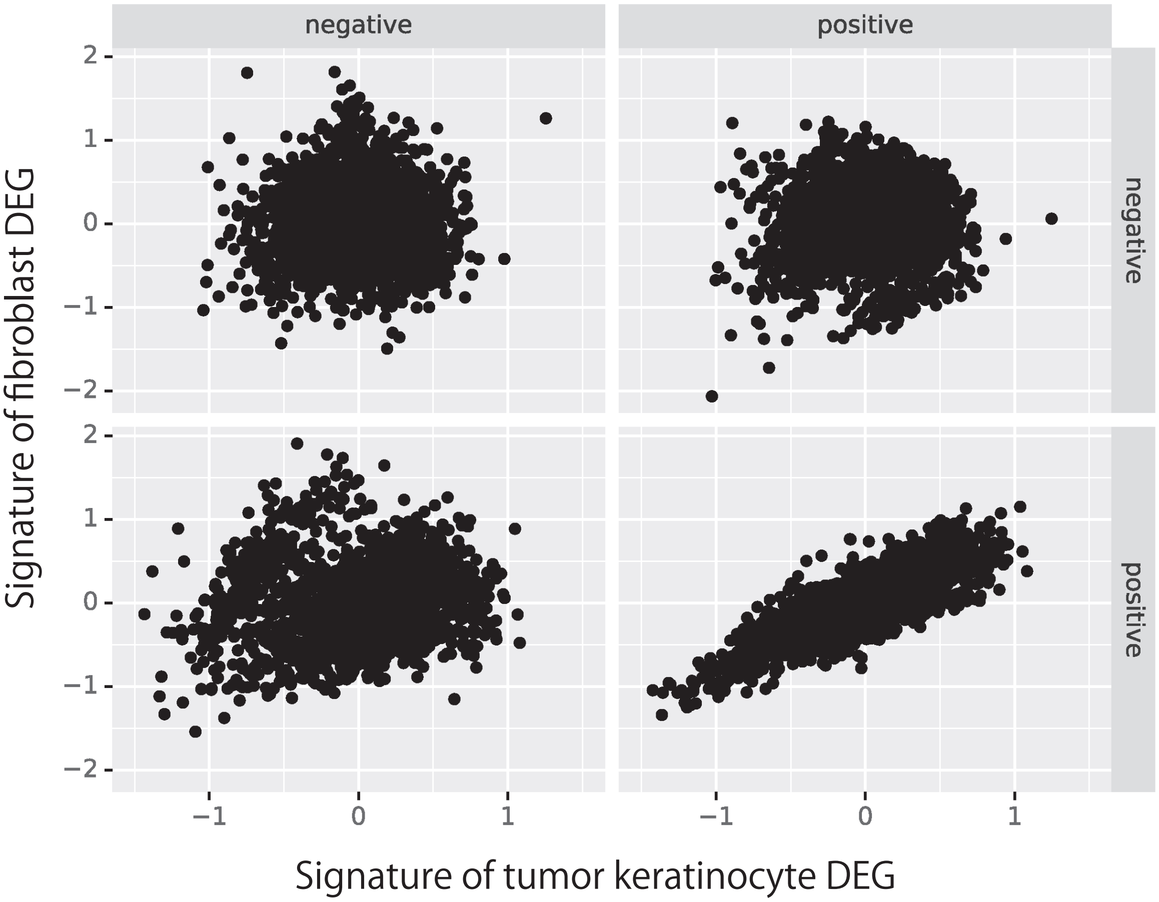


Figure 5: **TCGA analysis of colocalized fibroblasts and tumor keratinocytes** We calculated signature scores for positively and negatively regulated DEGs for fibroblasts and tumor keratinocytes in colocalization cluster 0 across SCC patients in TCGA dataset.

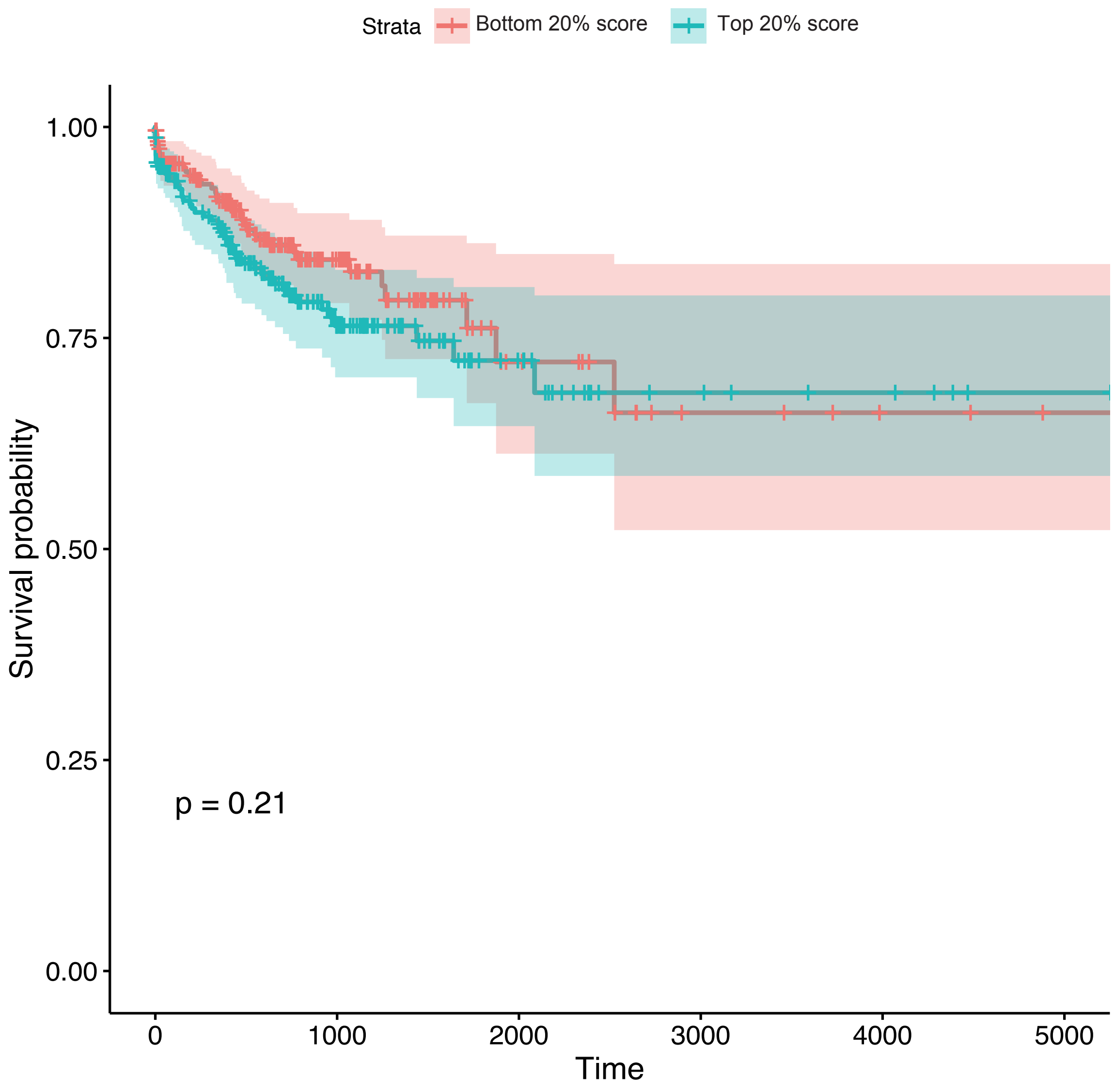


Figure 6: **Survival analysis for tumor keratinocyte signateres** Kaplan-Meier plot of the survival rate for patients with SCC with high and low signature scores of tumor keratinocytes belonging to colocalization cluster 0 (top and bottom 20%) in TCGA dataset.
